# A Tale of Two Chains: Geometries of a Chain Model and Protein Native State Structures

**DOI:** 10.1101/2024.02.10.579771

**Authors:** Tatjana Škrbić, Achille Giacometti, Trinh X. Hoang, Amos Maritan, Jayanth R. Banavar

## Abstract

Linear chain molecules play a central role in polymer physics with innumerable industrial applications. They are also ubiquitous constituents of living cells. Here we highlight the similarities and differences between two distinct ways of viewing a linear chain. We do this, on the one hand, through the lens of simulations for a standard polymer chain of tethered spheres at low and high temperatures and, on the other hand, through published experimental data on an important class of biopolymers, proteins. We present detailed analyses of their local and non-local structures as well as the maps of their closest contacts. We seek to reconcile the startlingly different behaviors of the two types of chains based on symmetry considerations.

## 1. Introduction

Polymer science [1-4], the study of chain molecules including linear polymers, is a flourishing subject that has led to life-changing progress in several technologies including plastics, textiles, and the design of novel materials. At the same time, linear chain molecules form the very basis of life including both the DNA molecule, whose information is translated into the sequence of amino acids, and proteins, which serve as amazing molecular machines in living cells. While conventional polymer models with stiffness have proved to be adequate for describing the relevant physics of the DNA molecule [5-7], the physical behavior of canonical polymers and proteins are strikingly different. In contrast to DNA, the structural changes during protein folding occur on multiple length scales at once, making it difficult to separate the relative contributions of the myriad interactions [8-13].

A linear chain is composed of many interacting monomers that are tethered together in a railway train topology. If the only interaction is self-avoidance, a single chain is in a coil phase whose large-scale behavior is in the same class as a self-avoiding walk. Upon adding an attractive interaction between pairs of non-adjacent monomers, the chain undergoes compaction into a highly degenerate compact phase at low temperatures [14-44]. While the notion of phases and phase transitions for a polymeric chain strictly refer to a chain with an infinite number of monomers, proteins are modest length chains, which yet exhibit several common characteristics. Notably, the compact state of proteins is modular and made up of two kinds of secondary building blocks, topologically one-dimensional helices [45], and two-dimensional sheets made up of zig-zag strands [46]. The helices and the strands are connected by turns or loops [47-49]. The nature of the ground states of compact polymers is qualitatively distinct from that of proteins and ordinarily do not exhibit any secondary motifs. The common characteristics of proteins are believed to derive from the shared backbone of distinct amino acid sequences.

Here we present an analysis of these two distinct classes of behaviors to understand their similarities and distinctions. We do this in two complementary ways. For conventional polymers, we study the simplest model of tethered hard spheres of diameter σ and a bond length equal to the sphere diameter. Following the standard nomenclature, we call this the tangent sphere model. We impose an attractive interaction between all pairs of non-adjacent monomers through a square well of range 1.6 σ and depth -1, which sets the energy scale without loss of generality.

A sphere is isotropic and looks the same when viewed from any direction. There is nevertheless a preferred axis at the location of each main chain sphere corresponding to the tangent along the chain or the direction along which the chain is oriented at that location. Replacing the spheres with objects such as unidirectional coins, uniaxial discs, allowing neighboring spheres to overlap, or adding side chains to the spheres along the main chain are all steps that break the spherical symmetry and yield ground state structures, which resemble protein structures to varying degrees [50-61]. To avoid clouding the issue by studying an approximate model for proteins, we resort instead to a careful analysis of experimental data of over 4,000 protein native state structures (see Materials and Methods). A side-by-side comparison of the local structures and non-local contacts of proteins to those of the tangent sphere model at both low and high temperatures provides a vivid picture of the different views of a chain molecule. Our goal here is to assess how well the tangent sphere model describes the protein backbone.

## 2. Materials and Methods

### 2.1. Our protein dataset

Our protein data set consists of 4,391 globular protein structures from the Protein Data Bank (PDB), a subset of Richardsons’ Top 8000 set [62] of high-resolution, quality-filtered protein chains (resolution < 2Å, 70% PDB homology level), that we further distilled out to exclude structures with missing backbone atoms, as well as amyloid-like structures (for full list of the PDB identifiers of protein structures in our database see Table S1 in the Supplementary Information of Refs. [58,63]). The program DSSP (CMBI version 2.0) [64] has been used to determine the backbone hydrogen bonding pattern and thus place each protein residue in context within a protein chain: within an α-helix, it is labeled an ‘α-residue’; within a β-strand, it is labeled a ‘β-residue’, or elsewhere, it is tagged as a ‘loop-residue’.

### 2.2. Numerical simulations of a chain of tethered tangent spheres

To obtain a set of independent equilibrium configurations of a chain of tethered tangent spheres comprised of N=80 spherical beads, subject to an attractive potential for a wide range of temperatures, we have employed standard replica exchange (RE) (or parallel tempering) canonical simulations [65,66]. Most of our simulations were carried out with a chain of 80 hard spheres of diameter σ. The model is a tangent sphere model because the bond length is also constrained to be equal to σ. At any temperature, the spheres do not self-intersect and are hard. We introduce a generic attractive square-well attraction of range R_att_ = 1.6σ, between all pairs of spheres, and magnitude ɛ, which sets the characteristic energy scale. The attractive interaction causes the chain to become compact at low temperatures.

The RE calculation [65,66] relies on a set of canonical simulations run in parallel at a set of M carefully chosen different temperatures, T_i_, i = 1, 2, · · · M. Each simulation represents a replica, or a system copy in thermal equilibrium. The key advantage is the possibility of swapping replicas at different temperatures without affecting the equilibrium condition at each temperature. This permits rapid equilibration even when there is a rugged free energy landscape. In each Monte Carlo (MC) simulation of one replica, new moves are accepted with the standard Metropolis acceptance probabilities [67]. We ensure that the number of swaps that entail exchange of replicas is large enough to ensure the fidelity of the statistics. The efficiency of the RE scheme depends on the number of replicas, the selected set of temperatures, as well as of the swap moves frequency. For best performance, the acceptance rate of swaps is tuned to be around 20% [68]. The RE simulations results are conveniently analyzed using the weighted histogram analysis method [69]. We employed 30 replicas, with a finer temperature mesh at lower values of the reduced temperatures (k_B_T/ɛ) in the range k_B_T/ɛ = 0.3-0.5 with a separation of neighboring temperatures of 0.02. In the k_B_T/ɛ = 0.5-1 interval, the separation of neighboring temperatures was 0.05, and for k_B_T/ɛ = 1-4, the separation interval was 0.2. We allowed for the RE swaps only between neighboring temperatures. The exchange moves were attempted every 100 MC steps per monomer. The length of the simulations was 10^9^ MC steps per monomer and per replica.

For sampling chain configurations at infinite temperature, we have used the standard Metropolis MC algorithm [67] in which all proposed updates of the chain configuration that respect its self-avoidance are accepted. In both simulation protocols, standard local moves, including crankshaft, reptation (or slithering-snake) moves, endpoint moves, and the nonlocal pivot move [70] are employed with equal probabilities.

The, N=80 beads, tangent sphere polymer exhibits two continuous (second order) ‘transitions’ (see inset of Figure 5(b)). At a temperature k_B_T/ɛ ∼ 3 (ɛ denotes the energy scale of attraction), there is a coil-to-globule transition (signaled by a kink in the specific heat per bead C_V_/Nk_B_). This is a finite-size counterpart of the θ-point for this system. At low temperatures k_B_T/ɛ ∼ 0.4 there is a second ‘transition’ into a compact globule phase, signaled by the maxima of the specific heat per bead C_V_/Nk_B_. The results we will present for the tangent sphere model are at infinite temperature and k_B_T/ɛ = 0.3.

In the next section, we will present some definitions and general observations pertaining to the local structure of a chain, especially in the continuum limit. We then go on to the Results Section, which is divided into two parts. First, we depict some similarities and some critical differences between the polymer model and protein structures in terms of power law behavior of certain geometrical measures of local structure. In the second part, we will highlight some striking differences between the geometries of the model polymer and protein native state structures. We then conclude with a brief discussion.

## 3. General considerations

### Three-body interactions and the self-avoidance of a continuum tube

We will begin our analysis with some definitions and general observations. For fixed bond length, a local chain conformation is specified by two angles θ and μ. θ is a measure of bond bending, a straight conformation has θ=π. The angle between successive binormals, μ, is the second angular coordinate and is the dihedral or torsional angle [63].

The standard model of a linear chain in polymer science, represented by tethered spheres, does not lend itself, in a natural manner, to housing helices, which are recurrent motifs in biomolecules [71-74]. In contrast, a tube, such as a garden hose, which can be thought of as a chain of discs, can be wound readily into a helix. The protein α-helix has a geometry akin to that of a tube wound tightly into a space-filling helix [75].

Consider a collection of uniform untethered hard spheres. The standard prescription for ensuring that spheres do not overlap is to ensure that the distance between the centers of every pair of spheres is no smaller than the sphere diameter. Pairwise interactions often capture the essence of interacting systems. The simplest generalization to a chain topology is tethered hard spheres (with the same self-avoidance constraint), which, as we have noted, is the simplest conventional model of polymer physics. In contrast, the alternative description of a chain as a tube leads to an unusual condition of self-avoidance. It has been shown that the correct way to determine whether a tube *in the continuum limit* is self-avoiding or not entails discarding pairwise interactions and invoking appropriate many-body interactions [75,76]. This is illustrated by considering the self-avoidance of a tube of non-zero thickness (see Figure 1). Knowledge of the distance between a pair of points on the tube axis (say A and B or B and D) does not discriminate between the two contexts of nearby points along the axis or in different parts along a chain. In the continuum limit, points A, B, and C, locally positioned along the axis, can become infinitesimally close to each other. This cannot be the case, however, for non-local points, where self-avoidance is a prime consideration, and one must ensure that the two points do not approach each other too closely. Knowledge of the coordinates of a pair of points does not inform you about the context that the two points are in – are they points locally along the axis (which can of course be arbitrarily close to each other) or are they non-local points (that may have come too close to each other possibly signaling an intersection)? This inability to discriminate between local and non-local pairs of points is at the root of the problem.

**Figure 1.**
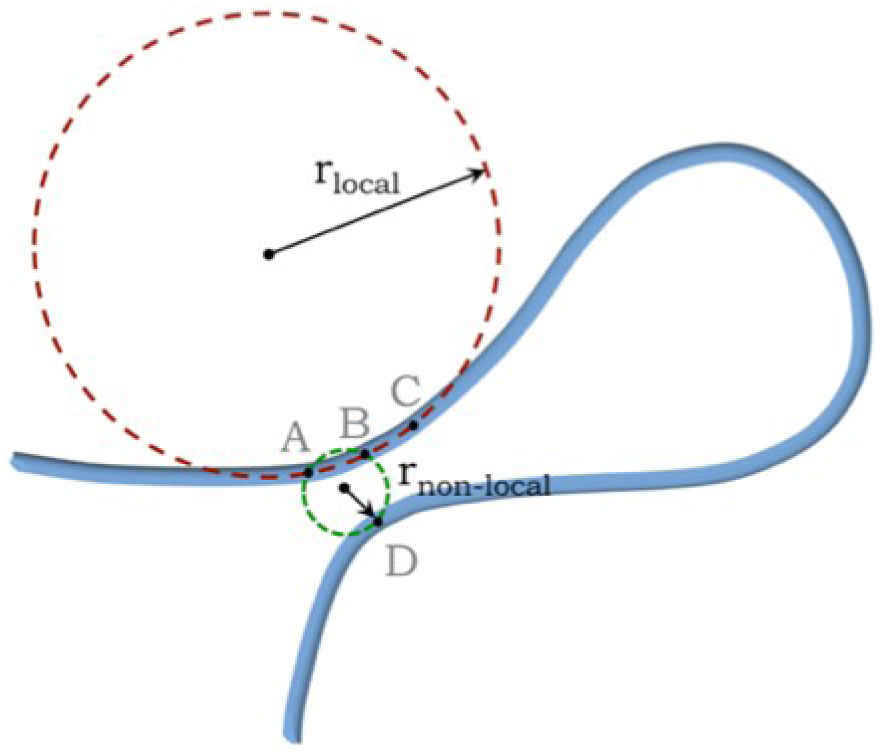
Sketch of the axis of a self-avoiding continuum tube (depicted in blue). The points A, B, and C lie alongside each other on the tube axis whereas point D is a nearby point from another part of the tube. The three-body prescription is to draw circles through all triplets of points on the tube axis and ensure that none of the radii is smaller than the tube radius. For a local triplet of points, one obtains the local radius of curvature whereas the non-local radius is a measure of the distance of closest approach of two parts of the tube.

The standard method in polymer physics of ensuring self-avoidance in the continuum limit is to first make the tube infinitesimally thin (a natural limiting case for a continuum chain of tethered spheres) and then use a singular δ-function potential [77] interaction: there is no energy cost as long as two points on the axis do not overlap exactly but there is an infinite energy cost when there is in fact an overlap. There are at least three problems with this [78]. First, the δ-function potential is singular (unlike say a familiar Lennard-Jones potential) and one needs to use renormalization group theory to introduce an artificial cut-off length scale to carry out the calculations and then demonstrate that this length scale can safely approach zero and yet retain the validity of the results. Second, the δ-function pairwise potential does not preserve the topology of a closed string – the number of knots is not necessarily conserved. Finally, in standard polymer physics, there is no description of the self-avoidance of a continuum self-avoiding tube (or surface) of non-zero thickness.

These problems are deftly averted by discarding pairwise interactions and working with a suitable many-body potential. In the continuum limit, there is a simple geometrical condition [75] to ascertain both whether a tube (and by extension a surface) is self-avoiding non-locally and is not too tightly wound locally. One can draw a circle through any triplet of points along the tube axis and measure the radius. The prescription for self-avoidance of a continuum tube is to consider all possible triplets, local or otherwise, and ensure that every one of the three body radii is greater than or equal to the tube radius. The radius of the circle passing through the triplet of points (A,B,C) in Figure 1 is the local radius of curvature as the points approach each other and results in a kink in the tube when the radius of curvature becomes smaller than the tube radius. On the other hand, the radius of the circle drawn through (A,B,D) is a measure of the distance of approach of two parts of the tube and must not be smaller than the tube thickness in order to respect self-avoidance. Likewise, the self-avoidance of a surface or layer (or a sheet of paper) of non-zero thickness necessarily entails discarding both pairwise and three-body interactions and working with a suitable four-body potential. One considers all quartets of points on the symmetry plane of a surface and draws spheres through each quartet. A surface is self-avoiding if the sphere radius of each quartet (local or non-local) is larger than the thickness of the surface. We note that this many-body prescription is strictly needed only in the continuum limit [76].

For a discrete chain, such as the ones we focus on in this paper, the pair-wise distance between a local pair of points is the bond length and there is no singular behavior at the local level because of a natural cut-off length scale. Also, a minimum threshold of the non-local three body radius can readily be measured by directly accessing the non-local pair-wise distance. We will make use of these simplifications for a discrete chain in our analysis below. It is important to note that there are myriad local interactions, including hydrogen bonds, as well as entropic effects that will play a key role in governing the local behavior of a real chain. Despite the absence of an imperative need to invoke many body interactions for a discrete chain, the three-body and four-body radii do yield interesting information. An infinitely large three-body radius signals co-linearity of the three points (bond bending angle θ equal to 180°; θ=0° is excluded because of steric overlap), whereas an infinite four-body radius is associated with planarity of the quartet of points (μ equal to 0°, 180°, or -180°).

## 4. Results

### 4.1. Power law scaling

Power laws often signify scale invariance and are a signature of an absence of a characteristic scale [79]. A liquid-vapor system at its critical point exhibits critical opalescence. The system appears milky white because light of all wavelengths scatter from the droplets and bubbles of liquid and vapor of all sizes thoroughly interspersed among each other. Another example of a non-trivial power law is the fractal dimension of a self-avoiding walk in three dimensions of around 5/3 [2]. Here we discuss a somewhat trivial but surprising realization of ‘universal’ power law behavior arising in the statistics of the local conformation of a discrete chain molecule. We alert the reader that, unlike in critical phenomena, here there is neither any many-body emergent behavior nor the need to invoke a system in the thermodynamic limit.

The procedure that we follow is simple. For a set of three (four) points, one can readily draw a circle (sphere) passing through them. The center of a circle (sphere) is the point equally distant from all three (four) points and can be determined as a solution of a suitable system of linear equations. In the case of three points, the solution is simple, and the radius of the circle R is related to the area A of the triangle passing through the three points and its sides a, b, and c: R=abc/4A. For four points, we merely solve the equations on a computer to obtain the radius R of the sphere.

Our goal here is to measure the radii R associated with many realizations of these points and obtain cumulative probability distribution functions of the inverse radius X=1/R. Figure 2 shows plots of the cumulative distribution function (CDF) of the inverse radii (X=1/R) (the probability P (1/r < 1/R) as a function of R that a given radius r is larger than R). The circles (spheres) in question are drawn through three (four) points (chosen consecutively along a chain and randomly in some instances) in three dimensions.

**Figure 2.**
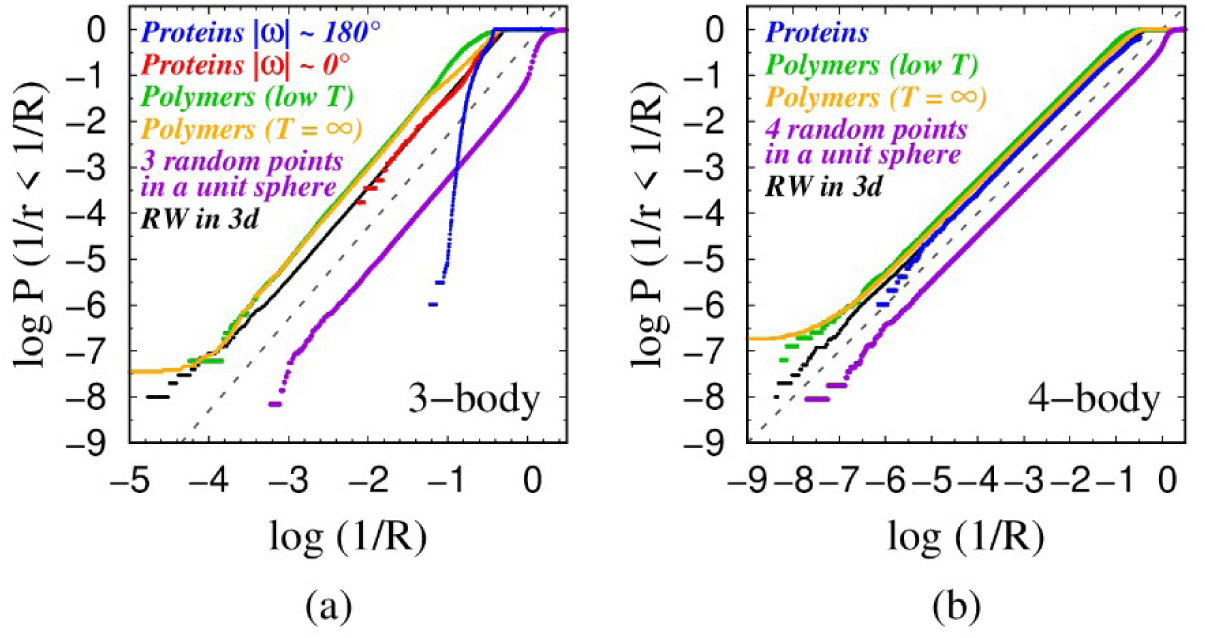
(a) Cumulative probability distributions of the inverse radii X=1/R of circles drawn through three consecutive points along different classes chains: (blue) 965,122 triplets of the backbones of globular proteins in our data set (defined by C_α_ atoms) when both the Ramachandran ω angles characterizing a consecutive triplet have canonical values of |ω| ≈ 180°; (red) 5,774 consecutive triplets of C_α_ atoms in globular proteins in which at least one of the two Ramachandran ω angles is|ω|≈0°; (green) 16,391,622 triplets taken from ≈200,000 low temperature (k_B_T/ε = 0.3) configurations, obtained using replica-exchange (RE) simulations, of a chain of 80 tangent spheres of diameter σ with an attractive square well potential of range R_att_ = 1.6σ ≈ 6Å; (orange) 25,598,976 triplets obtained from MC simulations at T = ∞; (purple) 143,557,206 triplets of points chosen uniformly from within a unit sphere in three dimensions; and (black) 100,000,000 two-step random walks in three dimensions with fixed bond length of 3.81Å. The gray dashed line has a slope of 2 and is a guide to the eye. (b) Cumulative probability distributions of the inverse radii X=1/R of spheres, whose surface passes through four consecutive points in different classes of chains: (blue) 957,723 local quartets along the backbones of globular proteins (employing C_α_ atoms); (green) 16,181,473 local quartets selected from ≈200,000 chain configurations, obtained from replica-exchange (RE) simulations, of a chain of 80 tangent spheres of diameter σ subject to an attractive square-well potential of range R_att_ = 1.6σ ≈ 6Å in the low temperature phase (k_B_T/ε = 0.3); (orange) 25,270,784 local quartets obtained from MC simulations at T = ∞; (purple) 112,754,340 quartets of points chosen uniformly within a unit sphere in three dimensions; and (black) 100,000,000 three-step random walks in three dimensions with a fixed bond length of 3.81Å. The gray dashed line is a guide to the eye and has a slope of 1. In all simulations, the bond lengths have been chosen to be 3.81Å equal to the mean value of the distance between the two consecutive C_α_ atoms along the protein chain. The distinctive behaviors of the purple curves (corresponding to the random points cases) occur because one can obtain circles of arbitrarily small radii, a situation precluded in the other cases due to steric considerations. The behaviors of the tangent polymer model at high and low temperatures are essentially the same. The local behavior is governed by the same steric constraints in both cases and the CDF does not change. In contrast, for real polymers, recent experimental studies [81,82] have shown the importance of the mechanical properties in determining the local curvature in the context of super-lubricity at the single molecule level.

We have studied: (a) consecutive triplets (quartets) of C_α_ atoms along the backbones of globular proteins when the Ramachandran ω angles characterizing a consecutive triplet have canonical values of |ω| ∼ 180°. Case (a) occurs in around 99.7% of the cases in globular proteins yielding the *trans* isomeric conformation of a peptide backbone, where the two neighboring C_α_ atoms are on opposite sides of the peptide bond, with a bond length approximately equal to 3.81Å [63]; (b) consecutive triplets (quartets) of C_α_ atoms in globular proteins in which at least one of the two Ramachandran ω angles has a rare non-canonical value of |ω|≈ 0° that occurs in ∼0.3% of cases. This happens when two neighboring C_α_ atoms are on the same side of the peptide bond, resulting in a shorter bond length of around ∼2.95Å. [63]. This corresponds to the so-called *cis*-conformation of a protein backbone [80]. Cases (a) and (b) are combined for the quartets because they show very similar behavior; (c) three (four) points selected from a two-step (three-step) self-avoiding random walk of a tangent sphere model (hard spheres of diameter 3.81Å and bond length 3.81Å with no other interaction besides the non-overlapping of hard spheres (polymer at infinite temperature); (d) three (four) points selected from a two-step (three-step) self-avoiding random walk of a tangent sphere model (spheres of diameter 3.81Å and bond length 3.81Å) subject to an attractive square-well interaction of range R_att_ = 1.6σ ≈ 6Å (polymer at low temperature); (e) three (four) points chosen randomly within a three-dimensional sphere of unit radius; and (f) three (four) points selected as points on a two-step (three-step) random walk (no self-avoidance or steric constraints) in three dimensions with a fixed bond length of 3.81Å (corresponding to the distance between consecutive C_α_ atoms in proteins [63]).

In the case of triplets (‘3-body’ case) shown in Figure 2(a), we see that the last five systems (b)-(f) exhibit power law behavior with approximately the same exponent of 2, but the canonical protein triplet does *not* (case (a) in Figure 2(a)).

For a canonical protein backbone in its *trans* conformation, quantum chemistry does not allow the bond bending angle θ to be greater than ≈150°, thereby preventing too large a value of R. In contrast, for a non-canonical protein backbone in its *cis* conformation [80], two consecutive C_α_ atoms along the protein chain are much closer to one another and the backbone in many of these cases has PRO residues. This stiffens up the protein backbone with respect to the canonical case permitting large bond bending angles, θ, that can almost reach ≈180°.

The second panel in Figure 2, Figure 2(b), shows similar ‘universal’ power law behavior for the CDF in the case of quartets (‘4-body’ case) of the inverse radius (X=1/R) of a sphere for various cases, detailed in the caption, this time with an exponent 1. We provide a simple rationalization of these findings in the next section for a triplet of points.

### Rationalization of the power-law exponent

To illustrate the origin of the power law behavior, we present here a simple derivation of the probability distribution P(X=1/R) of the inverse radii of circles drawn through three points of a two-step d-dimensional random walk with a fixed bond length b, which provides a natural length scale. We do not present a similar derivation for the radius of sphere passing through four points because it is more complex and is best handled numerically (which is what we have done). The quantity Xb is a dimensionless quantity, which enters in the derivation below. The input is p(θ), the probability distribution of the bond bending angle θ. For a d-dimensional random walk, the probability distribution p(θ) scales as [83],

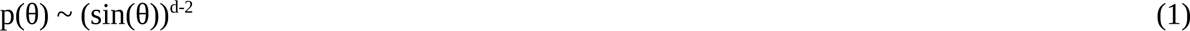

R and θ are related by

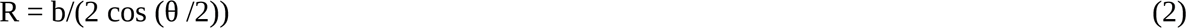

or equivalently:

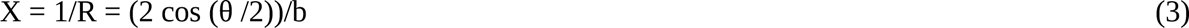

Noting that:

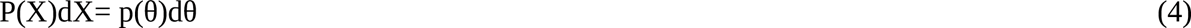

one obtains:

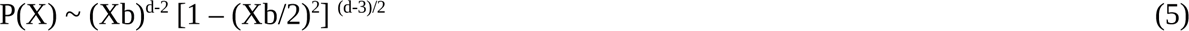

One thus obtains asymptotically (when Xb<<1 or in the large radius R limit) a power law behavior of the probability distribution P(1/R) with an exponent (d-2) for d>2 with a power law correction. This means that the cumulative probability distribution P(1/r < 1/R), being an integral of the probability P(1/R), displays a power law behavior with an exponent (d-1) (=2 in three dimensions) with a power law correction. This correction is relatively small when (Xb)^2^ is much smaller than 1 and yields good power law behavior as observed in Figure 2(a) and Figure 3(a). The pivotal quantity that determines the asymptotic exponent is the behavior of p(θ) when θ approaches 180° and the three points become co-linear. The difference in behavior in 2 and 3 dimensions is shown in Figure 3(b). The numerical simulations are in good accord with the prediction of Eq. (1).

**Figure 3.**
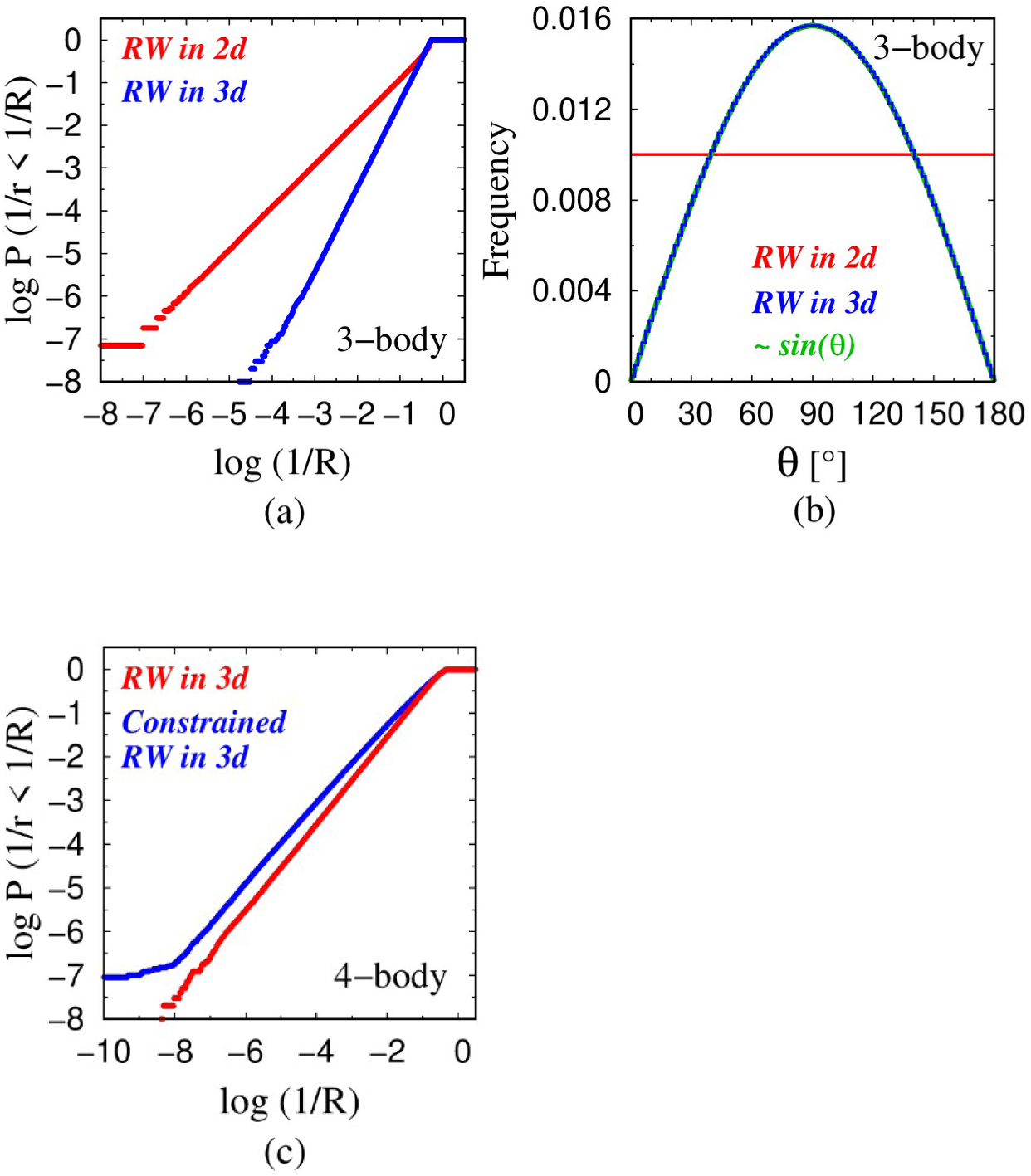
(a) Cumulative probability distributions of the inverse radii X=1/R of the circles drawn through three points of a two-step random walk with fixed bond length of 3.81Å, when the walk is performed in two dimensions (red) and in three dimensions (blue). (b) The distribution of the bond bending angle θ in the two-step random walk in two dimensions is uniform p(θ) = const. (red histogram), while, in three dimensions, p(θ) = sin θ (blue histogram and the green line). (c) Cumulative probability distributions of the inverse radii X= 1/R of the spheres drawn through four points of a three-step random walk with fixed bond length of 3.81Å, when the walk is performed in three dimensions (red) and for a constrained random walk. The constraint arises because the first three points (first two steps of the random walk) are sampled from a plane (sampling in two dimensions – after all, any three points do lie in a plane), while the final step (fourth point) is sampled in full three-dimensional space (blue). The constrained random walk corresponds to a fractal dimension 2 < d < 3 for the effective sampling of the variables controlling 1/R and reduces the steepness of the power law.

The ‘4-body’ case works in a similar manner to the ‘3-body’ case, except that the radius now depends on two independent variables. An unexpected sensitivity of the power law exponent to the choice of the 4 points is demonstrated in Figure 3(c). Quartets derived from a plain 3-step random walk in three dimensions (3d) exhibit power law behavior with an exponent of 1 (in accord with the results shown in Figure 2(b)). We have also considered a simple variant of the plain random walk that we call a constrained random walk. Here we define the first two points to lie along the x-axis. We then place the third point randomly on a pre-determined x-y plane. Superficially, this may not seem to be an onerous constraint, because any three points will necessarily lie in a plane. Nevertheless, a constrained random walk still shows power law behavior but with a distinct exponent, behaving as though it is in a fractal dimension regime. This is because the sampling of phase space is now altered in a relevant manner. We note that this regime may occur when a polymer system happens to be in the vicinity of a surface or a solid wall.

### 4.2 Chain geometries

We begin with an analysis of distinct local chain geometries. We compare the behaviors of the tangent sphere model at low and high temperatures, on one hand, and that of the native states of globular proteins, on the other. The local structures of these cases are shown in Figure 4 through their characteristic (θ,μ) plots [63]. These plots are drawn by measuring local pairs of bond-bending and dihedral angles for nearly a million monomers (for the polymer models) and residues (for protein native state structures). In contrast to the plots of model polymers (Figures 4(a) and (b)), the (θ,μ) plot for globular protein native state structures (Figure 4(c)) exhibits significant structure (distinct from the features present in the low temperature tangent sphere model) and signal the presence of the building blocks of helices and sheets.

**Figure 4.**
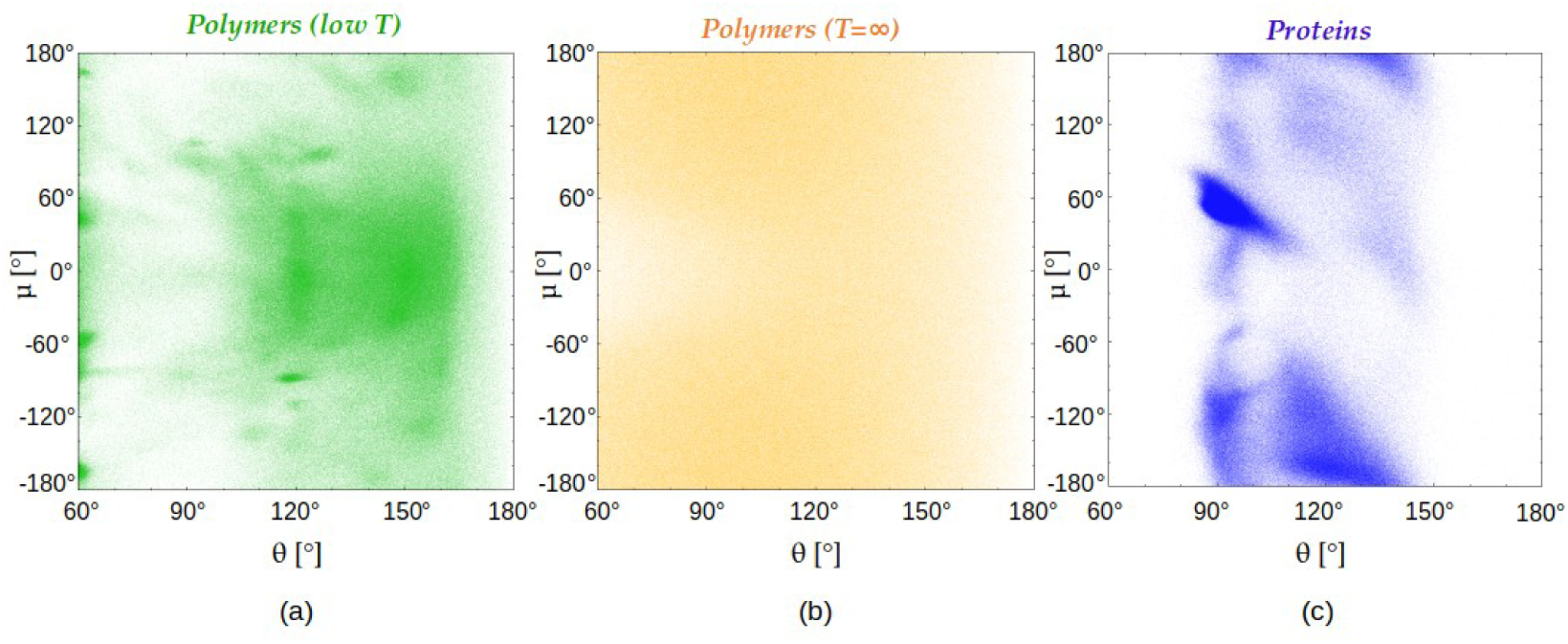
(θ,μ) cross plots of the bond bending angle θ versus dihedral angle μ for 966,505 randomly chosen monomers belonging to two different polymer classes and for the same number of residues in protein native state structures. (a) (light green) Polymer chain consisting of 80 tangent spheres of diameter σ subject to an attractive square well potential of range R _att_ = 1.6σ ≈ 6Å at low temperature (k_B_T/ε = 0.3) studied using RE simulations. (b) (orange) Polymer chain consisting of 80 tangent spheres of diameter σ at infinite temperature accessed by means of MC simulations with the only interaction being steric avoidance of all pairs of spheres. (c) (blue) For 966,505 residues of the 4,391 globular proteins in our data set.

The trends in Figure 4 are analyzed in Figure 5, which depicts a histogram of how far the nearest non-local monomer or residue is along the chain sequence. A non-local contact is defined to be one that is separated by at least three positions along the chain with the two beads nearest to each other (i,j) satisfying |i-j| ≥ 3. For both helices and turns, there are sharp maxima of close-by neighbors at a sequence separation of 3.

**Figure 5.**
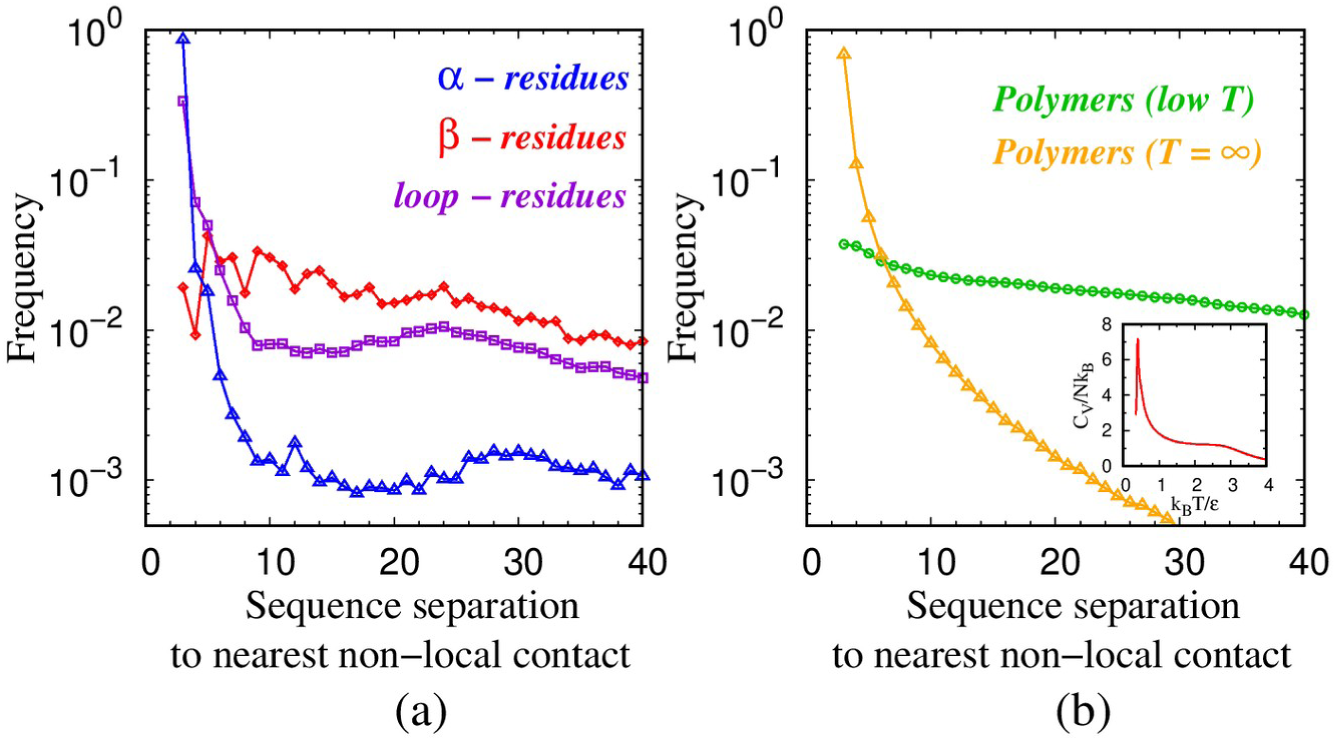
Frequency distribution of the sequence separation |i-j| along the chain, of the nearest non-local contact of bead i found at location j. In Panel (a), the blue points indicate 313,574 α-residues out of a total of 975,287 residues in our data set from 4,391 globular proteins; the red points are 214,501 β-beads; and the purple points denote an analysis of 442,821 loop-residues. (b) the green and yellow points show the distinct smooth behaviors of a tangent polymer model at low (k_B_T/ɛ = 0.3) and infinite temperatures. The behavior is monotonic with the closest non-local contacts always being close along the sequence. The inset shows the specific heat per bead C_V_/Nk_B_ for a 80 beads long tangent sphere polymer as a func_t_ion of the reduced temperature k_B_T/ɛ. There are two continuous (second order) ‘transitions’: at a high temperature ∼3, there is a coil-to-globule transition and, at ∼0.4, there is a transition into a compact globule.

There are at least two characteristic length scales associated with the conformation of a chain. One is the local behavior as measured by the local radius of curvature of a triplet of contiguous points. The second is the distance to the nearest non-local contact. For the space-filling conformation of a continuum tube of non-zero thickness, these two length scales become equal [75]. Figure 6 shows a histogram of the two scales, local and non-local, for several situations. The local radius is R = b/(2 cos(θ/2)), where θ is the bond bending angle associated with a local triplet. Structures in the histogram of local radii denote a preference for certain angles of θ. The α-residues exhibit a sharp peak corresponding to θ_α_≈92°, the β residues for θ_β_≈120°, and the loop residues have a pronounced maximum close to the helical value and a less prominent maximum around 111°, see Figure 6(a). These peaks are also reflected in the high-density regions in the (θ,μ) cross plot of protein native state structures, shown in Figure 4c. Figure 6b shows the histograms of the relevant non-local length scales for all five cases. All five curves exhibit a single peak denoting a relevant non-local length scale. Table 1 is a compilation of these characteristic local and non-local length scales.

**Figure 6.**
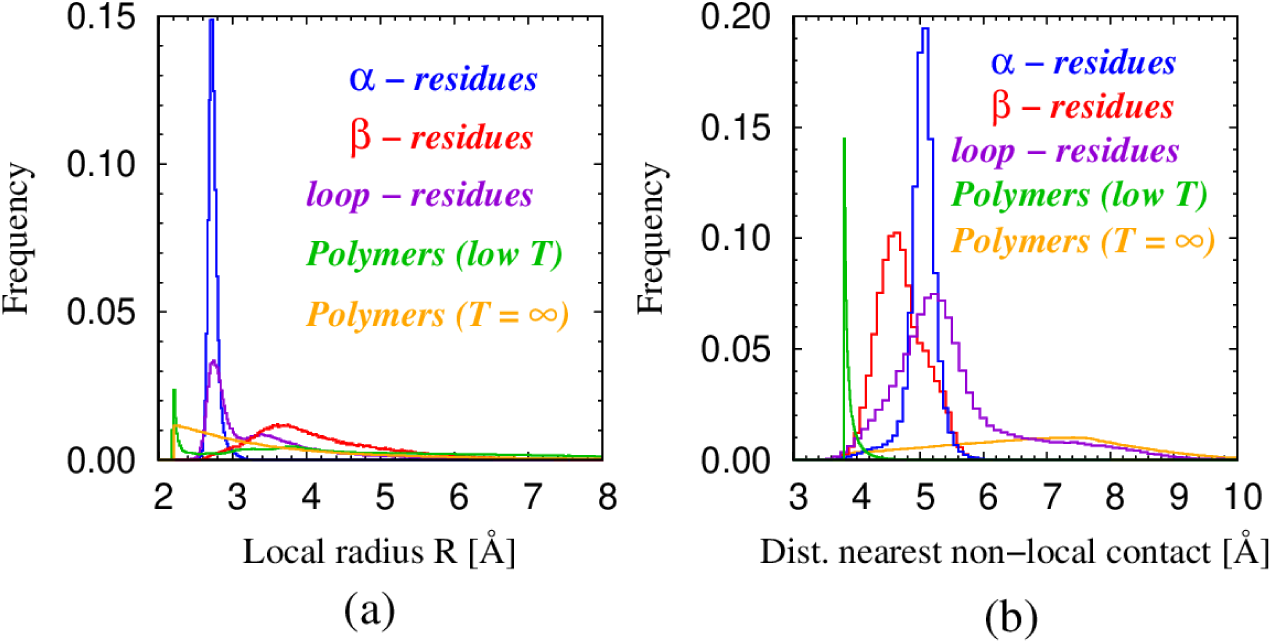
(a) Frequency distribution of the local radius of curvature R for five different classes of consecutive triplets. We consider only pure protein triplets in which all three residues are in the *same* structural class. There are 256,154 α triplets (blue) comprising ∼ 26% of all 970,896 triplets. There are 134,643 β triplets (red) and 313,923 loop triplets (purple). There are 16,391,622 low temperature polymer triplets (green) and 25,270,784 infinite temperature triplets (orange). (b) Frequency distributions of the distances to the closest non-local contact, defined as |j-i| ≥ 3, of monomers belonging to different classes, with the same color code as in panel (a).

**Table 1.** Characteristic local and non-local length scales for three residue types in proteins and for monomers of a tangent polymer chain in the low and high temperature phases.

| **Monomers** | **Most frequent value of local radius [Å]** | **Relevant non-local length scale [Å]** |
| --- | --- | --- |
| $\alpha$ – residues in proteins | 2.73 | 5.06 |
| $\beta$ – residues in proteins | 3.6 | 4.67 |
| loop – residues in proteins | 2.75 | 5.26 |
| Spheres in tangent polymers at low T | 2.21 | 3.81 |
| Spheres in tangent polymers at $T = \infty$ | 2.47 | 7.72 |

Figure 7 shows five representative chain conformations (denoted A-E), each having 80 monomers. We will present the key characteristics of these conformations to highlight similarities and especially the differences. Figure 7 also presents the contact maps for each of these five conformations to show for each monomer (labeled 1-80) the nearest, in distance, monomer (separated at least 3 positions along the chain) indicating in red, when the pairwise distance is greater than 6Å. Such distant contacts are rare in the polymer models unlike for the three protein chains. There is little structure in the globular polymer structure A. The infinite temperature polymer B is characterized by the nearest contacts being nearby in sequence, as evidenced by the points in its contact map being close to the diagonal.

**Figure 7.**
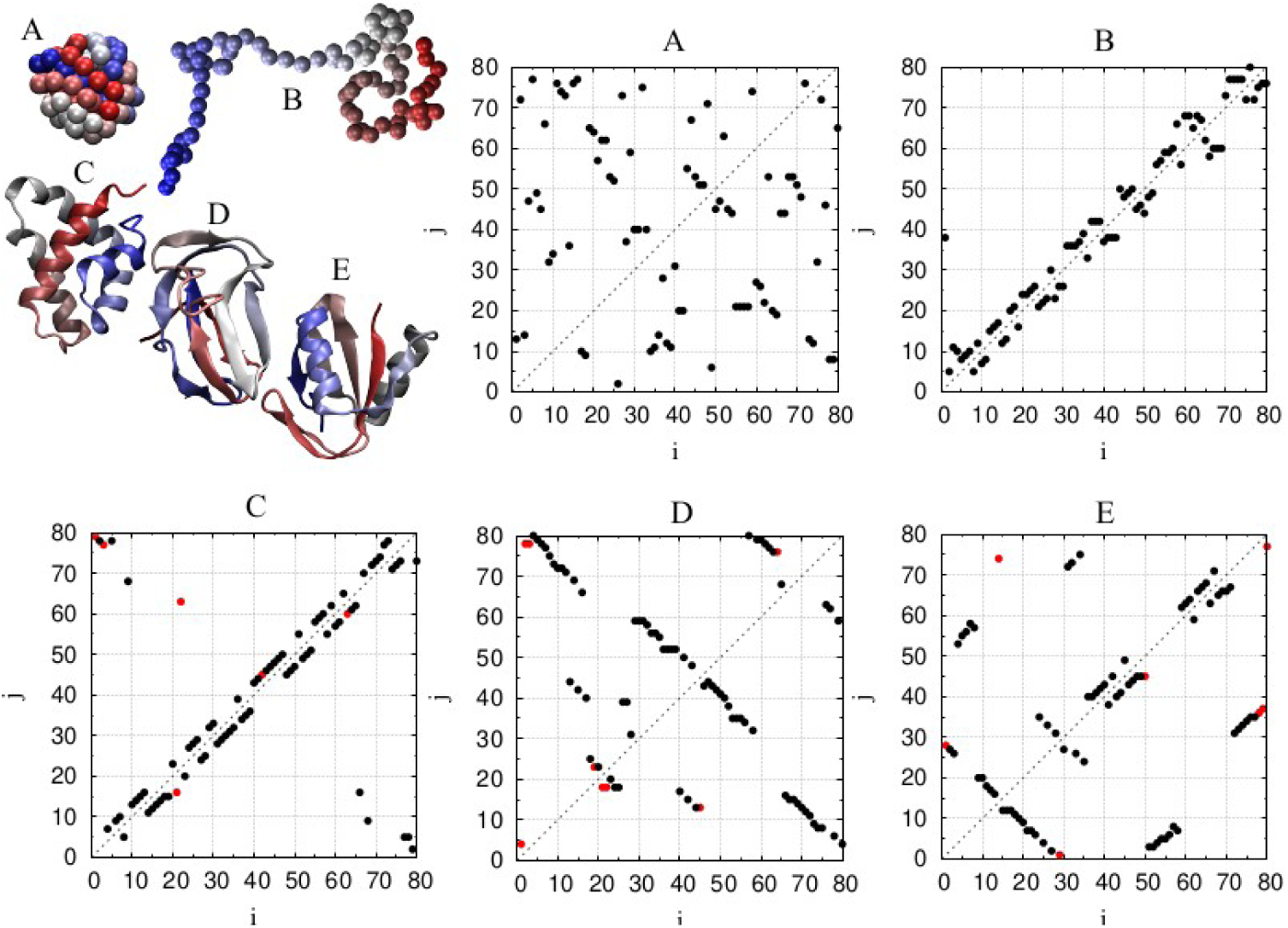
Representative conformations of chains of length 80: A) a low temperature globule and B) an infinite-temperature coil of a generic tangent polymer chain; C) all-α protein [PDB code: 3bqp, chain B]; D) all-β protein [PDB code: 1bdo, chain A]; and E) α/β protein [PDB code: 3l9, chain X]. The color coding of the conformation does not represent secondary motifs but rather depicts how far a monomer is along the sequence. The chain beginning is depicted in red color that morphs towards blue at the chain end. The contact maps shown in the other panels are those indicating the closest non-local contact j of a given monomer. The black points indicate those that are found within 6Å and the red points those that are found further away than 6Å, of these configurations. The choice of 6Å is dictated by the fact that the radial distribution function of proteins exhibits a pronounced minimum at this value [60].

This feature of locality of the closest contacts is also seen in protein α-helices C where coordinated closest contacts are of the (i,i+3) type. Furthermore, a distinctive pattern is seen for β sheets D, that display coordinated (i,j) contacts in which bead index j ≥ i+4 and takes on a coordinated pattern that is consistent with the situation of two strands coming together and forming parallel β sheets (when points in the contact map are parallel to the principal diagonal but shifted away from it) or anti-parallel β-sheets (when points in the contact map are placed along the directions that are perpendicular to the diagonal). The mixed α/β protein E has the features present in both α-helices and two types of β-sheets (parallel and anti-parallel). The similarity of the α helix contact map and that of the infinite temperature polymer is in accord with expectations that helices are more prone to nucleate from the coil phase than β sheets because of the prevalence of short-range contacts.

Figure 8 shows the relative importance of the local radius of curvature and the relevant non-local distance in determining the nature of compact conformations in the five cases. In all the five panels, we have scaled the quantities with the appropriate characteristic length scale defined in Table 1. Even though the scaling factors are clearly different, the behavior of the polymer chain is superficially similar at high and low temperatures. The scaled value of the closest non-local distance is substantially flat around a value of 1 with the fluctuations being a bit larger at infinite temperatures. In contrast, the local radius exhibits bigger swings in the low temperature conformation compared to that at infinite temperature. The α helix is special in that both the local and closest non-local scaled distances are close to each other and to the value 1. In the continuum limit, this equality would result in a space-filling helix [75]. The β regions and the loops do not exhibit this kind of behavior with the non-local scaled distance often being smaller than the scaled local radius of curvature.

**Figure 8.**
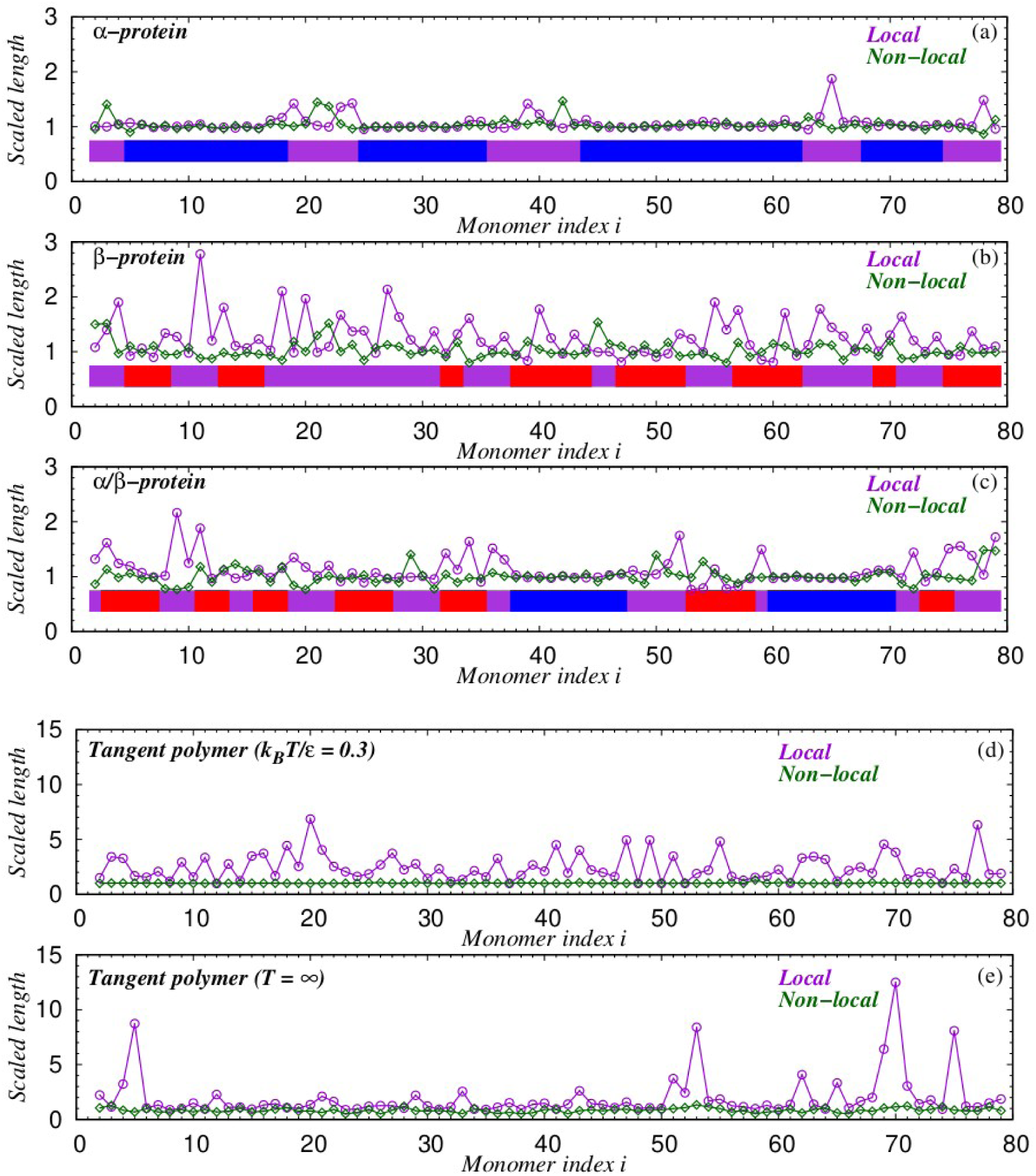
Scaled values of the local radius and minimal non-local distance for the five conformations shown in Figure 7. For each residue of the three proteins, depending on its type (‘α’, ‘β’, or ‘loop’) these quantities are appropriately scaled with the value of the corresponding characteristic length scale presented in Table 1. (a) all-α protein 80 residues long [PDB code: 3bqp, chain B]. (b) all-β protein 80 residues long [PDB code: 1bdo, chain A]. (c) α/β protein 80 residues long [PDB code: 3l9, chain X]. In the top three panels, the blue ribbons indicate α-helical parts of the sequence, the red ribbons indicate β-sheets, and the purple ribbons indicate loop regions; (d) a tangent polymer conformation at k_B_T/ɛ = 0.3; (e) a tangent polymer conformation at T = ∞.

## 5. Discussion

Our principal focus in this paper was to study the similarities and differences between chains viewed in two separate ways. The first, a baby model in polymer science, is a tangent sphere model subject to a square-well attraction. We have carried out extensive simulations of the model in the high and low temperature phases. We compare the behaviors with those of experimentally determined (and presented in the Protein Data Bank) structures of more than 4,000 proteins. The relevant local behavior can be measured by determining the radius of a sphere passing through a local triplet or the radius of a sphere, whose surface passes through a set of four consecutive monomers. Surprisingly, we found power law behavior of the probability distribution functions of the radii (with the notable exception of amino acid triplets in proteins). We presented a simple rationalization of this behavior.

We then went on to underscore the numerous distinctions between the model results and protein data. Proteins are complex molecules, which follow the rules of quantum chemistry [45,46,84,85] and are influenced by the interactions with the surrounding solvent molecules [86-97]. A protein is a distinct sequence of amino acids. Yet, proteins show remarkable common characteristics. They generally fold reproducibly and rapidly into their native state structures. These structures are directly implicated in protein function. The native state conformations are modular and made up of building blocks, notably helices and sheets comprised of zig-zag strands. These common characteristics arise because proteins share the same backbone despite having distinct sequences. An important open challenge is the determination of the simplest chain model that would suitably describe the striking features of the common backbones of protein native state structures.

## Author Contributions

Conceptualization, J.R.B, A.M., and T.Š.; methodology, J.R.B, A.M., and T.Š..; software, T.X.H. and T.Š.; validation, J.R.B, A.M., and T.Š.; formal analysis, T.Š.; investigation, J.R.B, A.M., T.Š.; resources, J.R.B, A.G., T.X.H., A.M., and T.Š.; data curation, T.Š.; writing—original draft preparation, J.R.B.; writing—review and editing, J.R.B, A.G., T.X.H., A.M., and T.Š.; visualization, J.R.B, A.M., and T.Š.; supervision, J.R.B; project administration, J.R.B.; funding acquisition, J.R.B, A.G., T.X.H., A.M., and T.Š. All authors have read and agreed to the published version of the manuscript.

